# Nanoflow ion-pairing LC–MS for ultra-low-input polar metabolomics and isotope tracing

**DOI:** 10.64898/2026.06.03.729938

**Authors:** Abigail E. Ellis, Rahul Deshpande, Abigail Cook, Craig P. Dufresne, Melanie Bailey, Susan S. Bird, Ryan D. Sheldon

## Abstract

Low-input and single-cell metabolomics remain constrained by the poor retention of polar metabolites in conventional reversed-phase nanoflow LC–MS workflows. Here, we establish nanoflow tributylamine (TBA) ion-pairing LC–MS as a platform for ultra-low-input polar metabolomics and stable isotope tracing. By adapting an analytical-flow TBA ion-pairing method to the nanoflow scale, this workflow extends the sensitivity and inline concentration advantages of nanoflow chromatography to charged metabolites involved in central carbon metabolism. Using mouse liver metabolite extracts, we show that the nanoflow method preserves chromatographic retention and separation of chemically diverse metabolite classes, including adenine nucleotides, nucleotide cofactors, TCA cycle intermediates, acyl-CoAs, and bile acid isomers. Despite loading 20-fold less tissue-equivalent material on column, nanoflow LC–MS produced higher signal intensity than the analytical-flow method for many metabolites. Across representative compounds, the nanoflow workflow reduced the biomass required for detection by approximately 20- to >600-fold, with pronounced gains for low-abundance metabolites such as NADPH and acetyl-CoA. TBA ion-pairing also enabled trap-and-elute nanoflow analysis of retained polar metabolites from single-cell-equivalent inputs. ATP was detected from one cell equivalent using both full-scan and targeted parallel reaction monitoring acquisition, with targeted acquisition further increasing signal over blank. Finally, we applied the workflow to stable isotope tracing in uniformly labeled ^13^C-glucose-treated cells. ^13^C-labeled ATP isotopologues were detectable from single-cell-equivalent input, and targeted acquisition improved isotopologue measurement near the detection limit. Together, these results demonstrate that nanoflow TBA ion-pairing LC–MS enables retained, high-sensitivity analysis of polar metabolites from ultra-low inputs and provides a foundation for extending central carbon metabolite analysis and isotope tracing toward single-cell-scale applications.

## INTRODUCTION

Single-cell and low-input molecular profiling has transformed biological measurement by shifting analysis from population averages to distributions of cellular states. Single-cell RNA sequencing has made it practical to profile thousands of individual cells, revealing transcriptionally distinct subpopulations and asynchronous differentiation programs that are obscured in bulk measurements ^1, 2^. However, RNA abundance provides only an indirect view of protein activity and metabolic phenotype. Mass spectrometry (MS)-based methods extend low-input analysis to proteins, lipids, and metabolites, providing molecular measurements that are complementary to sequencing-based approaches ^3, 4^. Because metabolites are direct substrates and products of biochemical pathways, low-input metabolomics has the potential to provide a functional readout of cellular state that is not accessible from transcript abundance alone.

Progress in MS-based low-input analysis has been most rapid in proteomics and lipidomics, where analytes are generally compatible with reversed-phase nanoflow liquid chromatography– mass spectrometry (nanoLC-MS). Miniaturized LC separations reduce chromatographic dilution, improve electrospray ionization efficiency, and maximize ion transfer from the source to the analyzer, thereby increasing sensitivity for sample-limited analyses. These advantages have enabled single-cell and low-input proteomics workflows, including isobaric mass tags, nanodroplet sample processing, and optimized low-nanoflow LC-MS configurations^3, 5, 6^. Similar gains have been reported in lipidomics, where nanoflow LC-MS and ion-mobility-enhanced workflows improve coverage and sensitivity from limited sample amounts ^7-9^. However, nanoflow LC-MS remains technically demanding, requiring careful management of sample loading, dead volume, column robustness, carryover, and clogging risk.

Adapting nanoflow LC-MS to metabolomics is particularly challenging because the metabolome spans a broader range of physicochemical properties than peptides or lipids. Central carbon metabolites, including organic acids, sugar phosphates, nucleotides, nucleotide cofactors, and acyl-CoAs, are often highly polar, ionic, and/or metal-sensitive. These properties lead to poor retention on conventional reversed-phase columns, early elution with matrix components, and limited separation of structurally related or isomeric metabolites ^10^. This is exacerbated on the nanoflow LC scale, where poorly retained compounds elute through the column during long sample loading times. Hydrophilic interaction liquid chromatography (HILIC) addresses many of these limitations at analytical flow rates (50-1000 µL•min^−1^) and has recently been adapted to the microflow scale (5 µL•min^−1^) ^11^, but their direct translation to nanoflow formats can be complicated by solvent compatibility, equilibration requirements, retention-time stability, and the increased vulnerability of nanoscale column to matrix contamination ^10, 12^.

Tributylamine (TBA) ion-pairing reversed-phase LC provides an alternative to HILIC for retaining anionic and highly polar metabolites while preserving compatibility with reversed-phase stationary phases and LC hardware. Protonated TBA associates with negatively charged metabolites, increasing their apparent hydrophobicity and promoting retention on reversed-phase columns. At analytical flow rates, TBA ion-pairing LC–MS has enabled sensitive measurement of metabolites from glycolysis, the pentose phosphate pathway, the TCA cycle, nucleotide metabolism, and CoA metabolism^12-17^. Unlike conventional reversed-phase LC, this approach substantially improves retention of polar metabolites while maintaining coverage of less polar and amphipathic compounds. However, TBA methods have historically been challenged by mobile-phase contaminants, ion suppression, isobaric interference, and metal-associated losses or signal instability. We recently developed a multi-stage purification and optimization strategy that reduces mobile-phase contaminants and metal ion interference, improving the analytical performance of TBA LC–MS and supporting its translation to nanoflow separations^18^. Adapting TBA ion-pairing LC to reversed-phase nanoflow platforms could therefore extend nanoflow sensitivity and inline concentration advantages to polar metabolomics in sample-limited applications.

Here, we describe a novel nano flow LC method using TBA ion-low-input polar metabolomics. This approach preserves broad chromatographic retention while substantially improving sensitivity over the analytical flow version of the LC method. We further demonstrate that TBA ion-pairing enables trap-and-elute nanoflow analysis of retained polar metabolites from single-cell-equivalent inputs, overcoming a key limitation of conventional reversed-phase nanoflow workflows in which polar analytes are lost during sample loading. Additionally, using uniformly labeled ^13^C-glucose tracing in cells, we show that ATP isotopologues can be detected and accurately quantified from single-cell-equivalent samples. Finally, the present manuscript provides the analytical foundation for an end-to-end single-cell metabolite profiling workflow demonstrated in an accompanying manuscript^11^. Together, these results establish TBA ion-pairing nanoflow LC-MS as a sensitive platform for low-input polar metabolomics and provide a foundation for extending central carbon metabolite analysis and isotope tracing toward increasingly sample-limited and single-cell-scale biological systems.

## RESULTS and DISCUSSION

To extend nanoflow LC-MS to low-input polar metabolomics, we adapted an established analytical-flow TBA ion-pairing method to the nanoflow scale. The goal of this translation was to preserve the broad retention of polar and charged metabolites provided by ion-pairing chromatography while leveraging the improved ionization efficiency and reduced sample requirements of nanoflow LC-MS. Nanoflow operation introduces distinct constraints, including large injection volumes relative to column flow, extended sample-loading times, and increased sensitivity to dead volume, so we first evaluated how method scaling affected sample introduction and chromatographic performance. We then compared nanoflow and analytical-flow methods using mouse liver metabolite extracts to determine whether nanoflow TBA ion-pairing LC-MS could maintain metabolite coverage while improving sensitivity for low-input analysis.

We adapted a well established analytical-flow (AF) TBA ion-pairing LC-MS method from our laboratory ^17-20^ to nanoflow (NF) conditions. The AF method uses a 150 mm × 2.1 mm, 1.8 µm Acquity Premier HSS T3 column operated at 250 µL/min with a 24 minute analytical gradient (Figure 1A). We scaled this method to a 150 mm × 75 µm, 2 µm PepMap column operated at 0.5 µL/min over a 15.2 minute method that included sample loading and analytical separation (Figure 1B).

**Figure 1.**
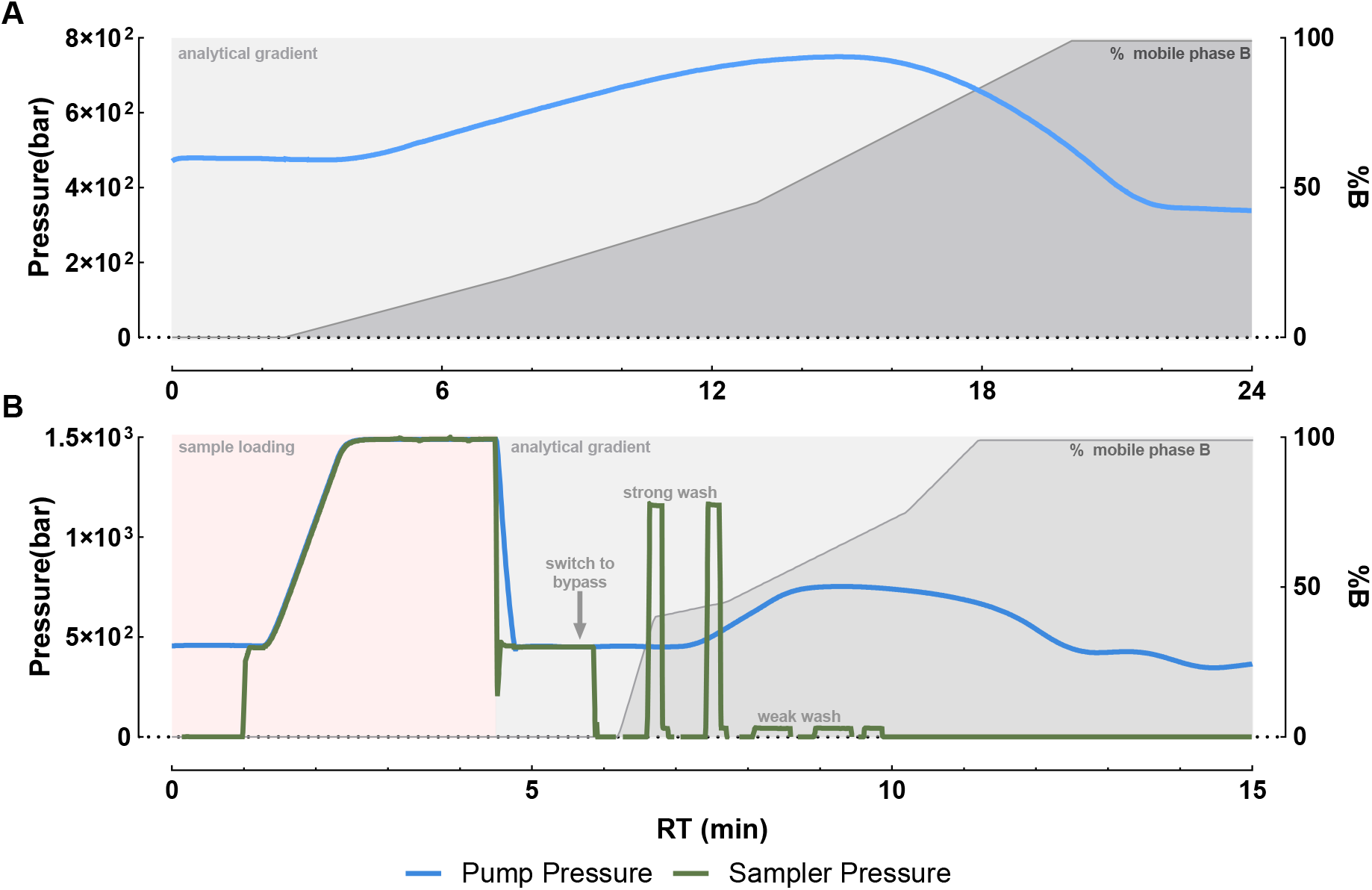
Method details for analytical and nano flow tributylamine liquid chromatography. (A) Pump pressure and LC gradient of the AF method. (B) Pump pressure, sampler pressure, and LC gradient of the NF method. Strong and weak washes are delivered by the autosampler metering device offline of the analytical flow path to prepare the sample loop for the next injection.

A key difference between AF and NF operation is the injection volume relative to the chromatographic flow rate. In the AF method, a 2 µL injection represents only 0.8% of the per-minute flow, allowing analytes to be introduced into the flow path as a compact sample plug. In contrast, under NF conditions, even a 0.1 µL injection corresponds to 20% of the per minute column flow. This relatively large injection volume can increase column pressure and disrupt elution profiles. To address this, a sample-loop injection is used to draw sample into a loop outside the analytical pump flow path. After the loop is loaded, the valve switches so that the analytical pump flow passes through the sample loop and carries the sample onto the column. This allows the sample to be delivered as part of the analytical flow rather than being pushed directly from the injector. However, this configuration requires additional loading time to account for dead volume in the sample loop and associated tubing. Here, sample loading required approximately 4.5 min. During this period, the flow rate was increased to maintain pressure just below the column maximum while transferring the sample through the loop and onto the column. The analytical gradient began at 4.5 min and continued for 10.7 min (Figure 1B).

TBA ion-pairing LC is particularly well suited for metabolite classes such as nucleotides, TCA cycle intermediates, and bile acids, which are effectively retained using this method. The ability to retain both highly polar compounds, such as nucleotides and TCA cycle intermediates, and more hydrophobic or amphipathic compounds, such as bile acids, highlights the mixed-mode strengths of this approach relative to conventional reversed-phase or HILIC methods. We therefore evaluated chromatographic performance of the nanoflow (NF) method relative to the established analytical-flow (AF) method using mouse liver metabolite extracts, which provide broad metabolic diversity and sufficient material for replicate analyses.

The adenine nucleotide series was well resolved by the NF method and eluted in the expected order of AMP, ADP, and ATP (Figure 2A), matching the elution order observed with the AF method (Figure 2B). Similarly excellent chromatographic behavior was observed for adenosine-containing cofactors, including NAD^⁺^, NADH, NADP^⁺^, NADPH, and FAD^⁺^, which were retained and separated using both methods (Figure 2C–D). FADH_2_ was not detected by either method, likely due to its low endogenous abundance and/or oxidation during sample preparation.

**Figure 2.**
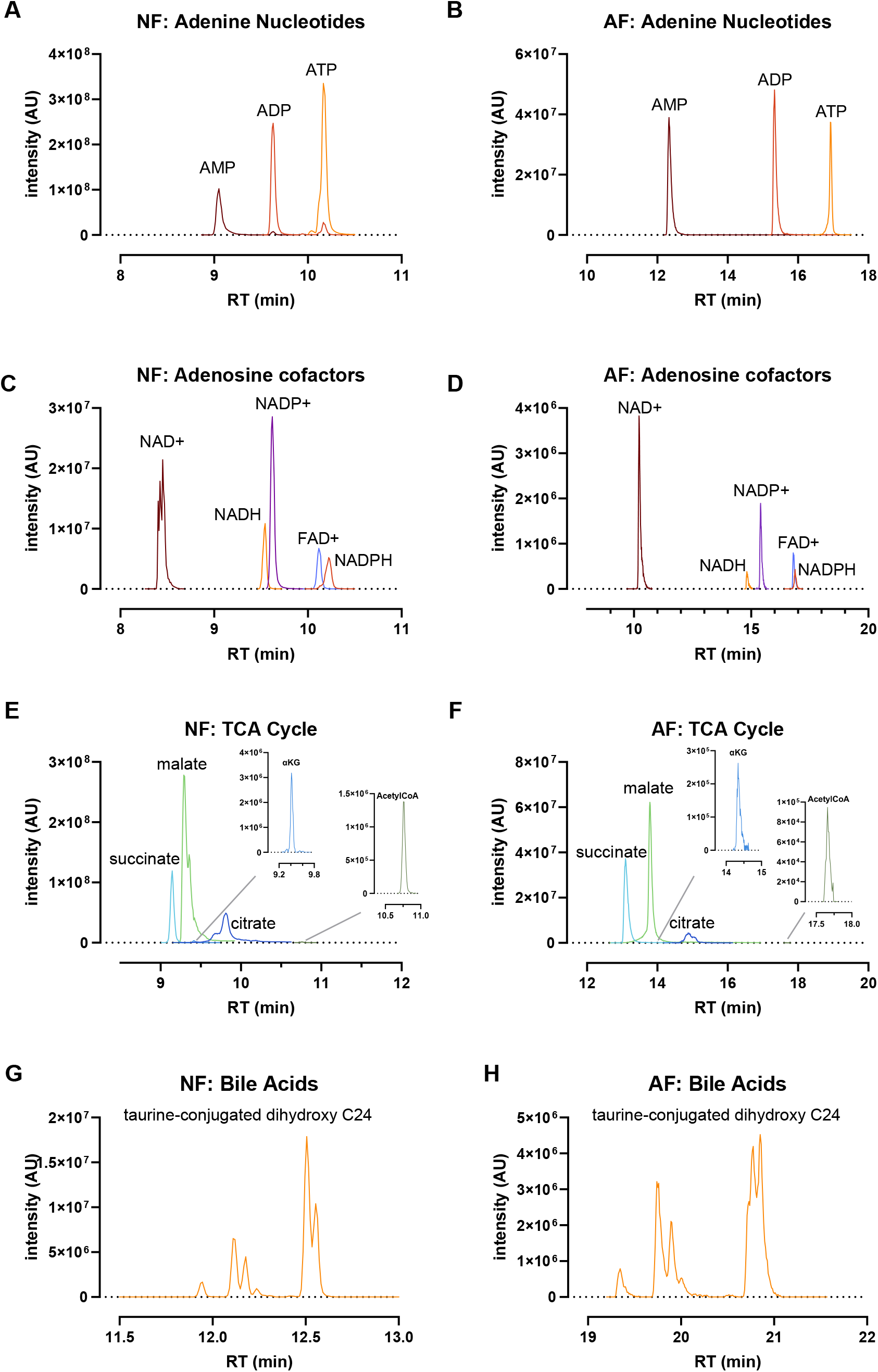
Comparative chromatographic performance using selected representative extracted ion chromatograms from NF (left) and AF (right) methods. (A-B), EICs of adenine nucleotides on the NF (A) and AF (B) methods. AMP, adenosine monophosphate; ADP, adenosine diphosphate; ATP, adenosine triphosphate. (C-D), EICs of adenosine containing cofactors from NF (C) and AF (D) methods. NAD+, oxidized nicotinamide adenine dinucleotide; NADP+, oxidized nicotinamide adenine dinucleotide phosphate; NADH, reduced nicotinamide adenine dinucleotide; NADPH, reduced nicotinamide adenine dinucleotide phosphate; FAD+, oxidized flavin-adenine dinucleotide. (E-F), TCA cycle intermediates on the NF (E) and AF (F) methods. αKG, alpha-ketoglutarate; CoA, coenzyme A. (G-H) EIC of taurine-conjugated C24 dihydroxy bile acids on the NF (G) and AF (H) methods. All chromatograms are representative of at least ten replicates.

TCA cycle intermediates and related central carbon metabolites are key readouts of energetic state and mitochondrial metabolism, but their highly polar, multianionic structures make them difficult to retain and resolve by LC–MS and can promote adsorptive losses on metal-containing surfaces in the LC flow path. The analytical-flow (AF) TBA ion-pairing method provided strong retention and resolution of these analytes, and this chromatographic performance was preserved after transfer to the nanoflow (NF) format (Figure 2E,F). Notably, despite lower biomass loaded on column, the NF method increased signal intensity for α-ketoglutarate and acetyl-CoA, two metabolites that are often challenging to measure due to low endogenous abundance.

Finally, we evaluated a taurine-conjugated dihydroxy C24 bile acid isomer group comprising five major murine species: taurodeoxycholic acid, taurochenodeoxycholic acid, tauroursodeoxycholic acid, taurohyodeoxycholic acid, and tauromurideoxycholic acid. These amphiphilic metabolites are poorly retained by HILIC, often requiring a complementary reversed-phase method for bile acid coverage. In contrast, the TBA ion-pairing method retained this isomer group in both AF and NF formats and resolved it into multiple discrete peaks, consistent with the presence of several prominent isomers in mouse liver extracts (Figure 2G,H). Together, these results demonstrate that transfer to the NF format preserves the broad chromatographic utility of the AF TBA ion-pairing method across chemically diverse metabolite classes.

A notable feature of the NF method was the substantial increase in signal intensity despite loading 20-fold less tissue equivalent on column relative to the AF method, 1.6 µg for NF compared with 32 µg for AF. Across the metabolite classes shown, signal intensity was approximately 5- to 30-fold higher with the NF method (Figure 2). This sensitivity gain was particularly apparent for lower-abundance or analytically challenging metabolites such as acetyl-CoA and α-ketoglutarate. In the AF method, these metabolites were only marginally detectable (Figure 2F), whereas in the NF method they produced clear peaks well above the 2 × 10^2^ intensity threshold used on this instrument (Figure 2E). These results indicate that adapting TBA ion-pairing chromatography to the nanoflow scale improves sensitivity while maintaining chromatographic separation of polar, charged, and amphipathic metabolites.

To quantify how NF LC-MS affected the amount of biomass required for metabolite detection, we prepared a half-log serial dilution of mouse liver metabolite extract beginning at 120 µg tissue equivalents/µL. Each dilution was analyzed using both the NF and analytical-flow (AF) methods, with injection volumes of 0.1 µL and 2.0 µL, respectively. To account for differences in background and absolute signal intensity between methods, peak areas were normalized to the corresponding blank and plotted as log_2_[signal/(blank + 1)]. Because blank signals were very low or absent for many metabolites, this transformation can produce relatively large values even near the detection limit. Therefore, metabolite detection was defined as a signal at least 3-fold above blank. Although the AF method produced higher absolute fold-over-blank values for some metabolites, the NF method maintained detectable signal at substantially lower on-column biomass.

Across the metabolites evaluated, the NF method reduced the amount of tissue required for detection in a compound-dependent manner. ATP was detectable with 16 ng of liver tissue equivalent on column using the NF method, compared with 316 ng using the AF method, corresponding to an approximately 20-fold reduction in required input (Figure 3A). This effect was even more pronounced for NADPH, which required only 1.6 ng on column with NF compared with 1012 ng with AF, a greater than 600-fold reduction (Figure 3B). Similarly, acetyl-CoA required 5.1 ng of tissue equivalent for detection by the NF method compared with 3200 ng by the AF method, also corresponding to a greater than 600-fold reduction in required input (Figure 3C).

Although absolute limits of detection depend on the mass spectrometer, ion source performance, and detector configuration, these comparisons were performed on the same MS system. Calibration and system-suitability standards were used to confirm comparable instrument performance across analyses. Therefore, while the specific detection limits and linear dynamic ranges may vary across platforms, these data demonstrate that adapting TBA ion-pairing LC-MS to the nanoflow scale provides a substantial sensitivity advantage for low-input metabolomics.

**Figure 3.**
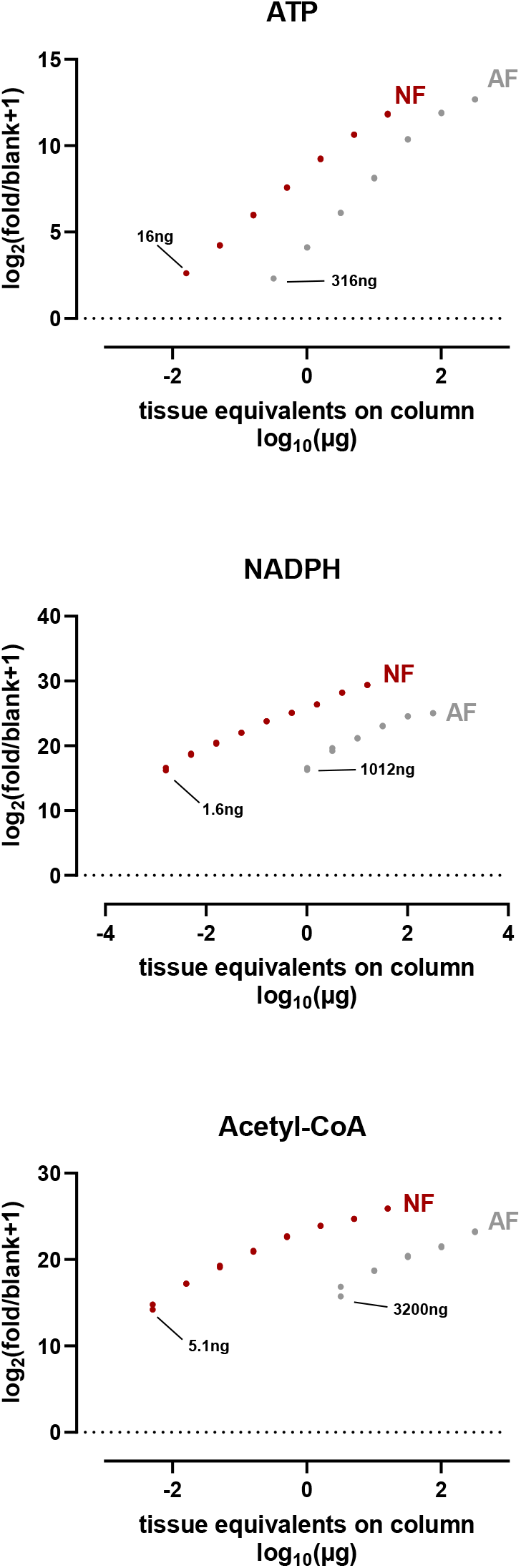
Nano flow ion pairing LCMS decreases on column biomass for metabolite detection. Mouse liver metabolite extracts were serially diluted and 0.1 and 2µL were injected on column using the NF and AF method, respectively. Shown are matrix dilution curves for (A) ATP, (B) NADPH, and (C) Acetyl-CoA. Only points with a fold-change over blank+1 greater than three are shown. Each curve was prepared in technical duplicate and analyzed separately. The lowest point detected at greater than 3-fold over blank +1 is annotated with the corresponding biomass on column.

Since polar metabolites are strongly retained on the NF method, we sought to evaluate whether we could increase loading volume to achieve a greater sample fraction on column. Using this approach, often referred to as trap-and-elute, retained analytes concentrate at the head of the column during loading and are subsequently eluted during the analytical gradient. In contrast, unretained compounds, such as highly polar metabolites in conventional reversed-phase workflows, will elute throughout the loading period and be lost before acquisition begins. This inline concentration is especially valuable for single-cell and other low-input applications, where maximizing sample transfer to the column is critical. We therefore evaluated whether nanoflow TBA ion-pairing LC-MS provided sufficient retention to support trap-and-elute analysis of polar metabolites at single-cell-equivalent input.

Metabolite extracts from a human B-cell line were diluted to one cell equivalent per 5 µL in mobile phase A without medronic acid. This volume is compatible with standard autosampler handling and approximates a practical volume for single-cell selection and processing without specialized liquid-handling systems. A 5 µL single-cell-equivalent sample was injected using the NF method and loaded onto the column over 20 minutes before initiation of the analytical gradient (Figure 4A). We selected ATP as a representative polar metabolite for evaluating trap-and-elute performance. In full-scan acquisition mode (70-850m/z), ATP, detected as the [M − H]^−^ ion at m/z = 505.9885, eluted at 23.87 min with a signal 3.4-fold above blank (Figure 4B). Because full-scan acquisition samples all ions across a broad m/z range, the ATP precursor represents only a small fraction of the accumulated ion population and is measured against substantial chemical background. We therefore evaluated targeted PRM acquisition, isolating ATP with a 1 Da window and using low collision energy (CE = 1) to transmit the intact precursor with minimal fragmentation. This approach increased ATP signal intensity to 10- to 20-fold above blank (Figure 4C). These results demonstrate that nanoflow TBA ion-pairing LC-MS can retain and concentrate polar metabolites from a single cell through extended sample loading and that targeted acquisition further improves detection near the We next asked whether this workflow could support stable isotope labeling analysis. Stable isotope tracing uses heavy-isotope-labeled metabolic precursors to measure isotope incorporation into downstream metabolites, providing insight into metabolic routing, substrate preference, and biosynthetic activity. This analysis is particularly challenging in low-input applications because metabolite signal is distributed across multiple isotopologues, reducing the signal intensity of any individual ion. To test this, we traced uniformly labeled ^13^C-glucose in a human B-cell line and evaluated ATP isotopologues at both the bulk-cell AF scale and the single-cell-equivalent NF scale. Nucleotides are particularly useful in stable isotope tracing experiments as labeling patterns can provide information on the activity of several upstream metabolic pathways. For example, glucose-derived carbon can be incorporated into ATP through several pathways, including labeling of the ribose moiety through the pentose phosphate pathway, which produces an M+5 isotopologue, and labeling of the adenine nucleobase through serine/glycine metabolism, one-carbon metabolism via 10-formyltetrahydrofolate, and/or CO_2_ fixation following TCA cycle oxidation.

**Figure 4.**
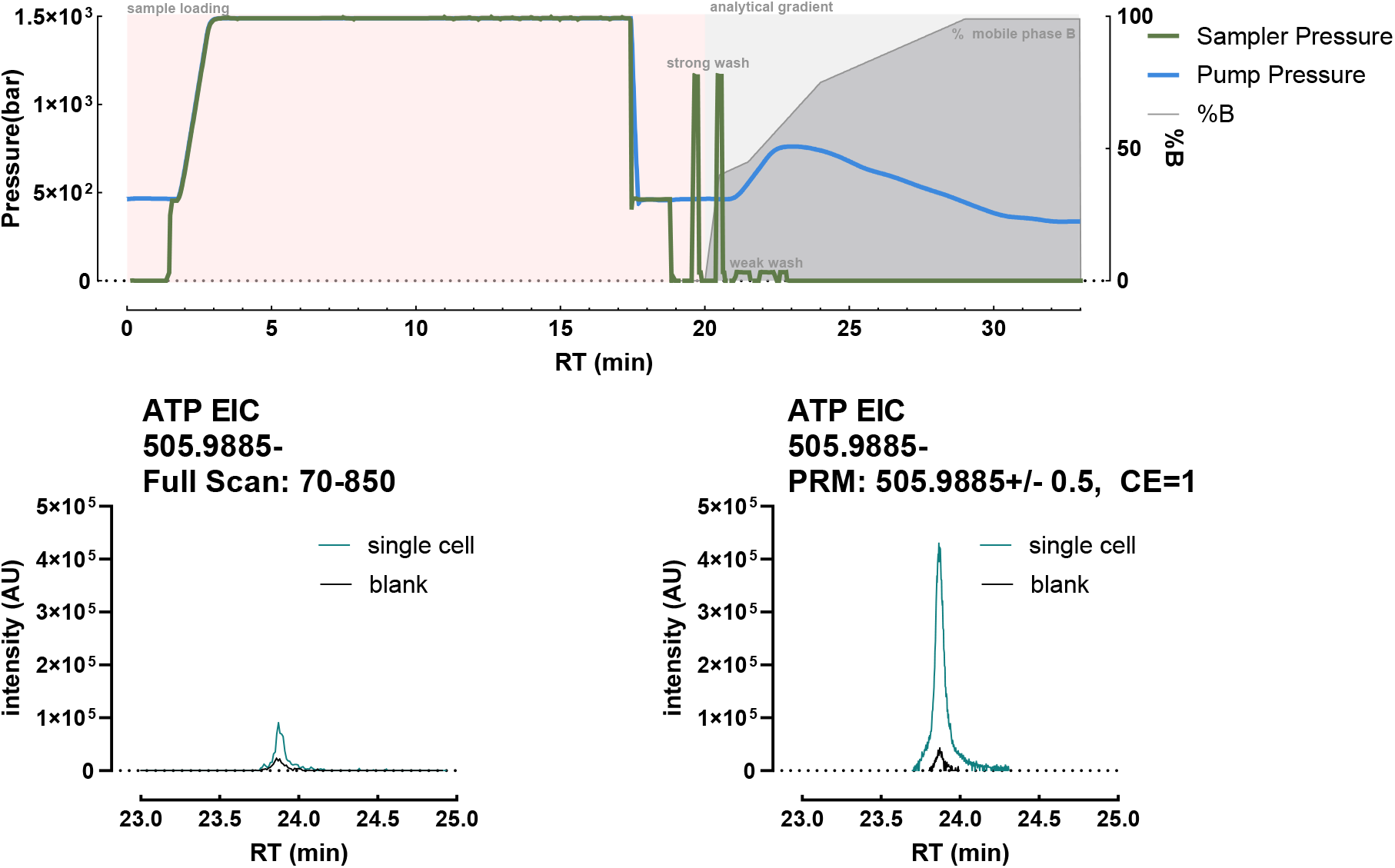
Single cell equivalent analysis of ATP. (A) A single cell equivalent metabolite extract in 5uL is loaded on the nano column over a 20 minute period, then eluted with the analytical gradient. Strong and weak washes are delivered by the autosampler metering device offline of the analytical flow path to prepare the sample loop for the next injection. (B) In full scan, ATP was modestly detected above the blank. (C) Targeted PRM acquisition mode boosts ATP signal intensity from a single cell equivalent injection versus the blank. EICs are representative of 9 replicates.

For AF analysis, 3 × 10^6^ cells were extracted, dried, and resuspended at a final concentration of 17,760 cell equivalents/µL. A 2 µL injection, corresponding to 35,520 cell equivalents on column, produced ATP isotopologues consistent with adenine labeling, observed as M+1 through M+4, and ribose labeling, observed as M+5 (Figure 5A). The same extracts were then diluted to one cell equivalent per 5 µL in mobile phase and analyzed using the NF method. Full-scan spectra showed clear ATP labeling across the M+1 through M+5 isotopologues at single-cell-equivalent input (Figure 5B). These labeled isotopologues were not observed in cells cultured with unlabeled ^12^C-glucose.

**Figure 5.**
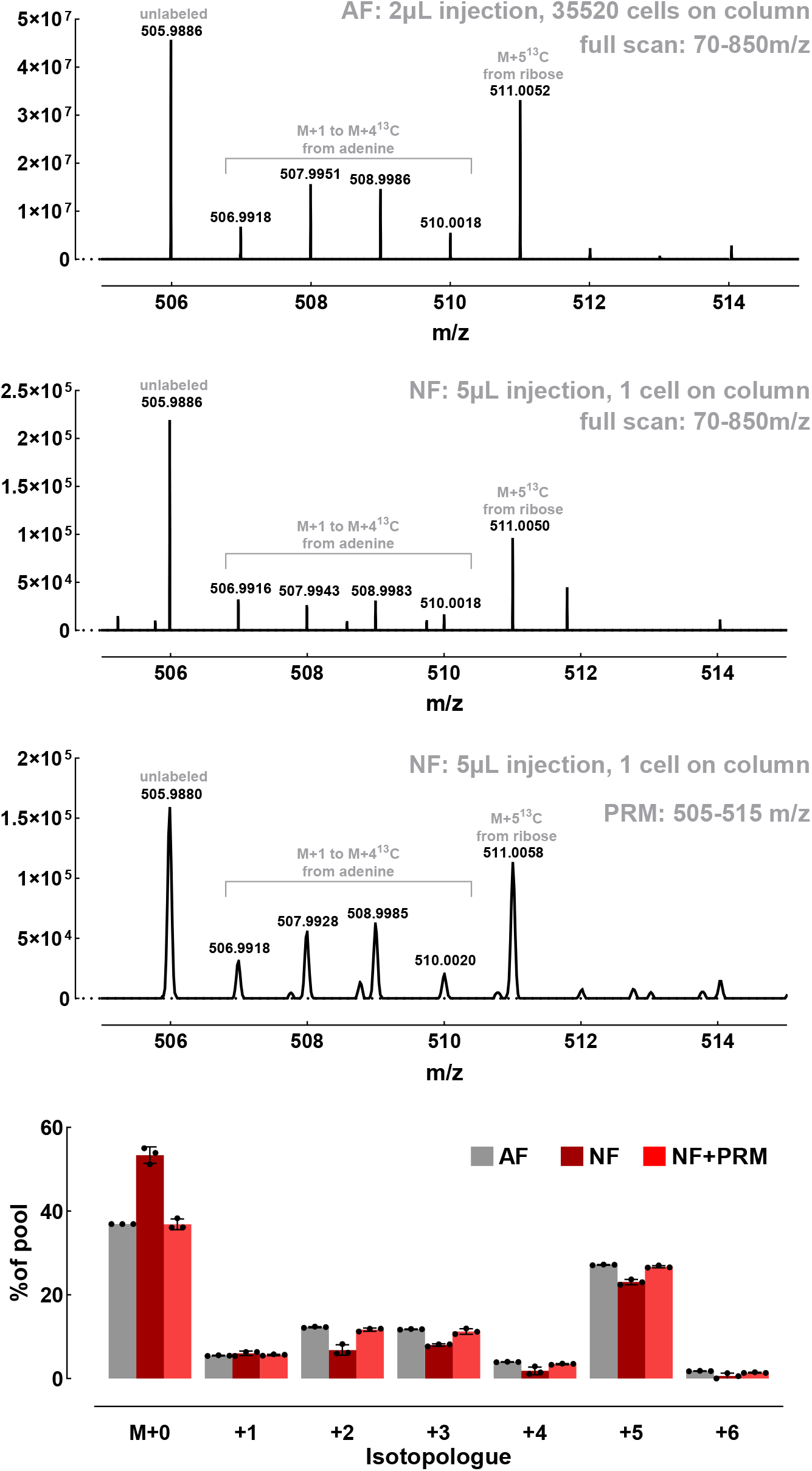
Stable isotope labeling of ATP from 13C glucose at the single-cell scale. (A) Full scan spectra of ATP at peak apex from 35520 cell equivalents on column using the AF method. (B) Full scan spectra of ATP at peak apex from 1 cell equivalent on column using the NF method. (C) Targeted PRM spectra of the ATP isotopologue envelope at peak apex from 1 cell equivalent on column using the NF method. (D) Each ATP isotopologue peak area expressed as a percent of the total ATP pool. Note, labeling was not observed in isotopologues 7-10 and are not shown here. Data represent n=3 biological replicates per condition.

Although ATP isotopologues were detectable by NF full-scan acquisition, the fractional abundance of the ^13^C labeled isotopologues was lower than observed with the AF method, suggesting reduced apparent isotopic fidelity near the detection limit. We reasoned that this reflected limited target ion transmission in full-scan acquisition. To test this, we reanalyzed the samples by PRM using a wide isolation window to capture the full ATP carbon isotopologue envelope and CE = 1 to transmit the unfragmented precursor in the MS2 channel (Figure 5C). This approach increased the measured fractional abundance of labeled isotopologues to levels comparable with the AF method (Figure 5D). Together, these data demonstrate that trap-and-elute nanoflow TBA ion-pairing LC-MS supports single-cell-equivalent detection of polar metabolites and can be extended to stable isotope tracing. Continued improvements in mass spectrometer sensitivity and targeted acquisition strategies should further improve quantitation of low-abundance isotopologues from metabolites successfully retained and delivered by this chromatographic workflow.

In this study, we introduce nanoflow TBA ion-pairing LC–MS as a new strategy for extending the sensitivity advantages of nanoflow chromatography to polar metabolomics. Although nanoflow LC–MS has reshaped sample-limited proteomics and lipidomics^3, 6, 8, 10, 12^, its impact in metabolomics has been constrained by a fundamental mismatch between conventional reversed-phase nanoflow separations and the polar, charged metabolites that define central carbon metabolism^13, 14^. By adapting TBA ion-pairing chromatography to the nanoflow scale, we address this limitation directly, enabling retained, trap-and-elute analysis of highly polar metabolite classes that are poorly suited to standard nanoflow reversed-phase workflows. This advance expands the chemical space accessible to nanoflow LC–MS beyond peptides, lipids, and hydrophobic metabolites to include organic acids, nucleotides, nucleotide cofactors, acyl-CoAs, and structurally related isomer groups. The resulting workflow combines broad polar metabolite coverage with the sensitivity and preconcentration benefits required for low-input analysis, providing a foundation for metabolomics in rare, spatially resolved, or otherwise sample-limited biological systems such as isolated cell populations, cerebrospinal fluid, laser-capture microdissection, fingerprint and sweat sampling, individual model organisms, and single-cell-scale applications.

A key advantage of this workflow is that retained polar metabolites can be concentrated during extended sample loading rather than lost before MS acquisition begins. This distinction is critical for low-input analyses, where sensitivity depends not only on improved ionization efficiency and ion transmission, but also on preserving as much of the available sample as possible and delivering it to the mass spectrometer as a focused chromatographic peak. The reduction in biomass required for metabolite detection, together with detection of ATP from a single-cell-equivalent input, demonstrates that nanoflow TBA ion-pairing can overcome a major limitation of conventional trap-and-elute nanoflow workflows for polar metabolomics.

Importantly, the platform also extends beyond metabolite detection to isotope-tracing measurements from single-cell-equivalent input. Detection of ^13^C-labeled ATP isotopologues after uniformly labeled ^13^C-glucose tracing shows that the method can recover pathway-informed metabolic information from extremely limited material. Because isotope labeling reports on substrate utilization, biosynthetic activity, and metabolic routing rather than abundance alone^21^, this capability provides a route toward functional readouts of pathway activity at extremely limited sample input levels.

Together, these findings establish nanoflow TBA ion-pairing LC–MS as a sensitive and chemically focused workflow for low-input polar metabolomics. By combining retention of central carbon metabolites with the preconcentration and sensitivity advantages of nanoflow LC–MS, this approach expands the metabolite classes accessible to sample-limited analysis. In doing so, it addresses a key gap in low-input and single-cell-scale molecular profiling: the ability to measure polar metabolites and isotope-labeling patterns that report directly on cellular metabolic state. This platform therefore provides a foundation for isotope-resolved measurements of metabolic pathway activity in rare, spatially resolved, or otherwise sample-limited biological systems. This analytical foundation is extended to end-to-end single-cell metabolite profiling in a companion manuscript^11^.

Limitations: TBA ion-pairing LC-MS generally requires a dedicated LC system because tributylamine is difficult to remove from the fluidic path and causes substantial ion suppression in positive-ion mode, making this workflow effectively limited to ESI-negative acquisition. In addition, although trap-and-elute loading enables concentration of retained analytes, unretained or weakly retained compounds can elute during the extended loading period and be lost before acquisition begins, a limitation common to nanoflow LC methods that use large sample volumes relative to flow rate. Throughput is also limited, with total run times of approximately 15–35 min per sample, largely constrained by sample loading time. Finally, LC-MS-based single-cell metabolomics is not currently compatible with the thousands-of-cells scale typical of droplet-based single-cell RNA sequencing, so experimental designs must be framed around focused questions that can be answered using relatively small numbers of cells. Thus, nanoflow TBA ion-pairing LC-MS is best viewed as a sensitive, chemically focused workflow for retained polar metabolites rather than a universal high-throughput single-cell metabolomics platform.

## EXPERIMENTAL SECTION

### Cell Culture

Human B-lymphoblastoid cells (GM24385, male; Coriell Institute) were cultured in Roswell Park Memorial Institute 1640 medium (RPMI 1640; 21870076, Gibco) supplemented with 15% fetal bovine serum (A5670701, Gibco), 1% penicillin-streptomycin, corresponding to 100 U/mL penicillin and 100 µg/mL streptomycin (15140122, Gibco), and glucose at a final concentration of 4 g/L (G7021, Sigma). Cells were maintained at 37 °C in a humidified incubator with 5% CO_2_.

### Stable Isotope Labeling with U-^13^C_6_-Glucose

For stable isotope tracing experiments, B-lymphoblastoid cells were cultured for 24 h in supplemented RPMI 1640 medium containing 22.9 mM uniformly labeled U-^13^C_6_-glucose (CLM-1396, Cambridge Isotope Laboratories). Unlabeled control cells were cultured in parallel using the same supplemented medium containing unlabeled ^12^C-glucose. After 24 h, cells were collected by centrifugation at 500 × g for 5 min. The medium was aspirated, and cell pellets were washed twice with ice-cold 0.9% saline (S8776, Sigma), flash-frozen, and stored at −80 °C until metabolite extraction.

### Mouse liver collection and processing

Four male C57B6/J mice were purchased from Jackson Laboratory and arrived at 12 weeks of age. They were housed for one week in a mouse facility on 12:12 light:dark cycle with free access to food and water. Mice were anesthetized with isoflurane and the liver was excised and snap frozen in liquid nitrogen. Frozen livers from each mouse were pooled together and cryo-pulverized to powder with a mortar and pestle. Pulverized tissue was maintained frozen at -80ºC until metabolite extraction. All animal procedures were approved by the Van Andel Institute IACUC.

### Metabolite Extraction

Metabolites were extracted using a modified Bligh-Dyer extraction procedure (chloroform:methanol:water, 2:2:1.8, v/v/v; PMID: 13671378). Liver tissue and cell pellets were extracted in ice-cold chloroform:methanol (1:1, v/v), with extraction volumes corresponding to 40 mg liver tissue per 1 mL solvent or 1.11 × 10^6^ cells per 1 mL solvent. After solvent addition, samples were vortexed for 10 s. Liver samples were homogenized for 30 s at 6 m/s using a liquid-nitrogen-chilled bead mill homogenizer. Cell extracts were not bead-milled. Both liver and cell extracts were then sonicated in a water bath for 5 min and incubated on wet ice for 30 min. Water was added to achieve a final chloroform:methanol:water ratio of 2:2:1.8.

For liver samples, tissue water content was assumed to be 70% and was accounted for when calculating the water addition. Samples were vortexed for 10 s, incubated on wet ice for 10 min, and centrifuged at 17,000 × g for 10 min at 4 °C. Following centrifugation, the lower organic phase, upper polar phase, and interphase protein pellet were separated. The upper polar phase was collected for metabolomics analysis.

For cell extracts, aqueous-phase volumes corresponding to 0.88 × 10^6^ cell equivalents were collected. For liver extracts, polar-phase volumes corresponding to 34 mg liver equivalents were collected and centrifuged a second time to remove residual particulates. After the second centrifugation, polar-phase volumes corresponding to 32 mg liver equivalents were collected. Cell and liver polar extracts were dried under vacuum.

### Sample Preparation for Nanoflow and Analytical-Flow LC-MS

Dried liver polar extracts were resuspended in LC-MS-grade water for liver metabolomics analyses. For chromatographic comparison between nanoflow (NF) and analytical-flow (AF) methods, liver extracts were prepared at 160µg tissue equivalents per mL, such that 0.1 µL injected by NF corresponded to 1.6µg tissue equivalent on column and 2.0 µL injected by AF corresponded to 32 µg tissue equivalent on column. For biomass dilution experiments, liver extracts were serially diluted in half-log increments. Each dilution was analyzed by both NF and AF methods using injection volumes of 0.1 µL and 2.0 µL, respectively. Tissue equivalents on column were calculated from the extract concentration and injection volume. Dried cell extracts used for single-cell-equivalent analyses were resuspended in mobile phase A without medronic acid. Extracts were diluted to one cell equivalent per 5 µL, and 5 µL was injected for single-cell-equivalent NF analysis.

### Nanoflow Tributylamine Ion-Pairing LC-MS

Nanoflow LC-MS analyses were performed using a Vanquish Neo liquid chromatography system coupled to an Orbitrap Exploris 240 mass spectrometer through an EASY-Spray nano-electrospray ionization source operated in negative-ion mode (Thermo Fisher Scientific). Mobile phase A consisted of LC-MS-grade water (W6, Fisher) containing 3% LC-MS-grade methanol (A456, Fisher). Mobile phase B consisted of LC-MS-grade methanol. Both mobile phases contained 10 mM tributylamine (90780, Sigma), 15 mM acetic acid (A11350, Fisher), and 0.1% medronic acid (v/v; 5191-4506, Agilent). Solvent calibration was performed using the Vanquish Neo “Calibrate Solvents” scripts to ensure accurate solvent proportioning with the ion-pairing mobile phases. The weak needle wash matched mobile phase A, and the strong needle wash consisted of 99% LC-MS-grade acetonitrile (A955, Fisher). The needle was washed after sample draw using 3 s of strong wash at 80 µL/s followed by 5 s of weak wash at 80 µL/s, with the default wash-cycle time.

Separations were performed on an EASY-Spray PepMap Neo C18 column, 2 µm, 75 µm × 150 mm (ES75150PN, Thermo Fisher Scientific). The column temperature was maintained at 35 °C, and the analytical flow rate was 0.5 µL/min. The column void volume was 0.444 µL, with a maximum pressure of 1500 bar. Samples were loaded using the loop-offline direct-injection configuration. Loading used combined flow/pressure control with a maximum flow of 15 µL/min and maximum pressure of 1500 bar. Injection volumes varied depending on the experiment and were 0.1, 1.0, or 5.0 µL. Because injection volume directly affected sample-loading time, the gradient was delayed at 100% mobile phase A until loading was complete. Loading times were 6.2, 7.0, and 20 min for 0.1, 1.0, and 5.0 µL injections, respectively. Total method times were 15.2, 16.0, and 33.0 min, respectively. After sample loading, the analytical gradient proceeded as follows: 0.5 min ramp to 40% B, 1.0 min ramp to 45% B, 2.5 min ramp to 75% B, 1.0 min ramp to 99% B, and 4.0 min hold at 99% B. Following the analytical gradient, the column was returned to 0% B and washed/equilibrated for 3 min. Prior to the next injection, the column was equilibrated using fast equilibration with pressure control set to 1200 bar and an equilibration factor of 3.

The mass spectrometer was set to acquire during the injection/loading period to monitor early-eluting or weakly retained compounds. Source parameters were as follows: static spray voltage, −1500 V; ion transfer tube temperature, 325 °C. Full-scan acquisition was performed from m/z 70–850 at a resolution of 240,000 with the RF lens set to 70%. The AGC target was set to standard, maximum injection time was 100 ms, and microscans were set to 1. For single-cell-equivalent detection of ATP, ATP was targeted as the [M − H]^−^ ion at m/z 505.9885 using a 1 Da isolation window and normalized collision energy of 1. For ATP isotopologue analysis, the isolation center was 510.9885 with an isolation window of 11 to capture the full carbon isotopologue envelope (505.9885-516.0220). Resolution was set to 11,250 FWHM.

### Analytical-Flow Tributylamine Ion-Pairing LC-MS

Analytical-flow LC-MS analyses were performed using a Vanquish Horizon liquid chromatography system coupled to an Orbitrap Exploris 240 mass spectrometer equipped with a heated electrospray ionization source operated in negative-ion mode (Thermo Fisher Scientific). Samples and standards were injected at 2 µL and separated using an Acquity Premier HSS T3 column, 1.8 µm, 2.1 mm × 150 mm (186009472, Waters), fitted with an Acquity Premier VanGuard cartridge, 1.8 µm, 2.1 mm × 5 mm (186009473, Waters). The column temperature was maintained at 35 °C, and the flow rate was 0.25 mL/min.

Mobile phase A consisted of LC-MS-grade water (W6, Fisher) containing 3% LC-MS-grade methanol (A456, Fisher). Mobile phase B consisted of LC-MS-grade methanol. Both mobile phases contained 10 mM tributylamine (90780, Sigma), 15 mM LC-MS-grade acetic acid (A11350, Fisher), and 0.01% medronic acid (v/v; 5191-4506, Agilent). The analytical gradient was 24 min in length and proceeded as follows: 0–2.5 min, 0% B; 2.5–7.5 min, 0–20% B; 7.5–13 min, 20–45% B; 13–20 min, 45–99% B; and 20–24 min, 99% B.

Between injections, a 16 min reverse-flow wash and re-equilibration method using a separate Vanquish Horizon pump was used to backflush the column. For the wash method, mobile phase A was unchanged, and mobile phase B consisted of 99% LC-MS-grade acetonitrile (A955, Fisher). The wash gradient was performed as follows: 0–3 min, 100% B at 0.25 mL/min; 3–3.5 min, 100% B with a ramp to 0.8 mL/min; 3.5–7.35 min, 100% B at 0.8 mL/min; 7.35–7.5 min, 100% B with a ramp to 0.6 mL/min; 7.5–8.25 min, 100–0% B with a ramp to 0.4 mL/min; 8.25–15.5 min, 0% B with a ramp to 0.25 mL/min; and 15.5–16 min, 0% B at 0.25 mL/min.

Mass spectrometer source parameters were as follows: spray voltage, −2500 V; sheath gas, 60 arbitrary units; auxiliary gas, 19 arbitrary units; sweep gas, 1 arbitrary unit; ion transfer tube temperature, 320 °C; and vaporizer temperature, 250 °C. Full-scan MS1 data were acquired in the Orbitrap over m/z 70–850 at a resolution of 240,000 with the RF lens set to 35%. Full-scan data were used for metabolite quantification and isotope-tracing analysis. Data-dependent MS/MS was also acquired for metabolite annotation using higher-energy collisional dissociation with stepped normalized collision energies of 15, 30, and 45. MS/MS spectra were acquired in the Orbitrap at a resolution of 15,000 with a 2 m/z isolation window, and data-dependent acquisition was limited to a maximum of five MS/MS scans per cycle.

### Data Processing and Analysis

Raw LC-MS files were processed using FreeStyle software (Thermo Scientific, 1.8.65.0) to general extracted ion chromatograms and Skyline (v. 26.1.0.057) for peak area quantitation. Extracted ion chromatograms were generated using a mass tolerance of 5ppm and retention-time windows determined from authentic standards. EIC data points were exported and plotted in GraphPad Prism. EICs shown are representative of at least 10 independent replicates.

Peak areas were integrated using automatic integration and inspected for consistency across dilution series and method comparisons. For comparisons between NF and AF methods, peak areas were compared using matched tissue-equivalent dilution series and corresponding solvent blanks. For biomass-limit experiments, peak areas were normalized to the corresponding blank and expressed as log_2_(fold over blank + 1). Detection was defined as a signal at least 3-fold above blank. The minimum biomass required for detection was assigned as the lowest tissue equivalent on column that met this criterion. Each dilution was prepared in technical duplicate and analyzed separately.

For stable isotope tracing experiments, ATP spectra were exported from FreeStyle and plotted in GraphPad Prism. Peak areas for each isotopologue were quantitated in Skyline using a 5ppm mass tolerance. Isotopologue fractional abundance was calculated as the peak area of each isotopologue divided by the summed peak area of the ATP M+0 through M+10 isotopologue envelope. For single-cell-equivalent isotope tracing, full-scan and PRM acquisition were compared to evaluate isotopologue detection near the limit of detection. These studies were completed using three biological replicates.

## ACKNOWLEDGMENTS

LC-MS analysis was completed in the Van Andel Institute Mass Spectrometry Core (RRID:SCR_024903) and supported financially by the VAI MeNu (<u>Me</u>tabolism and <u>Nu</u>trition) program (SCR_027494). R.D., C.P.D, and S.S.B. are employees of Thermo Scientific.

